# Feeling the force: how pollen tubes deal with obstacles

**DOI:** 10.1101/266106

**Authors:** Jan T. Burri, Hannes Vogler, Nino F. Läubli, Chengzhi Hu, Ueli Grossniklaus, Bradley J. Nelson

**Affiliations:** Multi-Scale Robotics Lab, Department of Mechanical and Process Engineering, ETH Zürich, 8092 Zürich, Switzerland; Department of Plant and Microbial Biology and Zürich-Basel Plant Science Center, University of Zürich, 8008 Zürich, Switzerland

## Abstract

**Highlight:** Pollen tubes literally feel their way through their environment to avoid obstacles as they deliver male gametes to the ovule. We measured their force sensitivity to understand this remarkable behavior.

**Abstract:** Physical forces are involved in the regulation of plant development and morphogenesis by translating mechanical stress into the modification of physiological processes, which, in turn, can affect cellular growth. Pollen tubes are tip-growing cells that provide an ideal system to study processes induced by exposure to mechanical stress. We combined a lab-on-a-chip device with cellular force microscopy to mimic and quantify the forces that are involved in pollen tube navigation upon confronting mechanical obstacles. Several stages of obstacle avoidance were identified, including force perception, growth adjustment, and penetration. We have experimentally determined the perceptive force, which is the force threshold at which the pollen tube senses the obstacle, for *Lilium longiflorum* and *Arabidopsis thaliana*. In addition, we provide evidence that pollen tubes are capable of penetrating narrow gaps by increasing turgor pressure. Taken together, our data indicate that pollen tubes sense physical barriers and actively adjust their growth behavior to overcome them.

## Introduction

The integration of physical stress, growth, and development has attracted burgeoning attention in plant developmental biology. Plant cells are able to respond to external physical forces in several ways, for example by directing cell wall synthesis to the site of increased stress, or by modifying biochemical and biophysical processes to regulate cell expansion (Hamant *et al*., 2008; Heisler *et al*., 2010; Ellinger and Voigt, 2014; Kesten *et al*., 2017). The ability to perceive mechanical forces is fundamental to all plant cells (Darwin and Darwin, 1880; Jaffe, 1973; Jaffe and Forbes, 1993; Braam, 2004; Chehab *et al.*, 2009). Until now, numerous studies have shown the effect of external mechanical forces on the growth patterns of plant organs (thigmomorphogenesis) with respect to cytoskeletal organization, cell wall integrity, biosynthesis, cellular anisotropy in expansion, and patterns of phytohormone responses (Lintilhac and Vesecky, 1981; Haley *et al.*, 1995; Yahraus *et al*., 1995; Wymer *et al*., 1996; Legué *et al*., 1997; Lynch and Lintilhac, 1997; Hamant *et al*., 2008; Haswell *et al*., 2008; Monshausen and Haswell, 2013).

Pollen, comprising a vegetative cell that encloses the two sperm cells, forms a cellular protrusion called the pollen tube (PT), which forms upon pollen hydration and germination. The PT grows through stylar and ovarian tissues to reach the ovules, where it bursts and releases the two sperm cells to effect double fertilization (Borg *et al*., 2009). The advancing apex of the elongating PT, driven by turgor pressure, generates forces large enough to penetrate through these tissues. During their growth, PTs need to perceive external mechanical obstacles and exert sufficient penetrative forces. Meanwhile, PTs need to reconfigure their growth behavior to withstand externally applied compressive stress. The PT with its fast growth rate and rapid response to external stimuli offers a perfect model system to investigate the impact of mechanical cues at the single-cell level (Chebli and Geitmann, 2007). The molecular mechanisms underlying the accurate navigation of the PT towards the ovule, by perceiving and responding to guidance cues, have been thoroughly investigated over the last decade. Numerous chemotropic factors have been identified, such as sugars, calcium, nitric oxide, lipids, adhesins, and secreted peptide signals serving as guidance cues (Hülskamp *et al*., 1995; Ray *et al*., 1997; Wolters-Arts *et al*., 1998; Mollet, 2000; Higashiyama *et al*., 2003; Prado *et al*., 2004; Sanati Nezhad *et al*., 2014; Higashiyama and Yang, 2017; Qu *et al*., 2015).

Until now, the vast majority of research on PT guidance has focused on the mediation of directional growth by biochemical factors. However, a major hurdle for the PT on its way to the ovule is the physical interaction with the surrounding tissues. The growing interest on the influence of mechanical forces acting as cues for cellular behavior and thereby regulating growth and morphogenesis, has triggered the demand for new techniques to study the effect of mechanical stimuli at the single-cell level. With the advances in microelectromechanical systems (MEMS) and microfluidics, various lab-on-a-chip (LOC) devices have been fabricated for the high-throughput analysis of PTs and their interactions with biochemical and mechanical cues (Horade *et al*., 2013; Agudelo *et al*., 2013; Sanati Nezhad *et al*., 2014; Shamsudhin *et al*., 2016; Yanagisawa *et al*., 2017; Hu *et al*., 2017*b*; Hu *et al*., 2017*a*). Despite the insights offered by these LOC devices, they still lack the ability to directly measure the variation of cytomechanical properties or the forces exerted on the PTs. Force values have to be extracted using modeling methods, such as finite element method (FEM)-based models, which has been done for example to determine the penetrative force of *Camellia japonica* PTs (Sanati Nezhad *et al*., 2013). These values are approximations and strongly depend on the accuracy of the model and input parameters. Most models cannot incorporate the complex nature of cytomechanical phenomena, and the trade-off between accuracy and available computation power makes it difficult to model dynamic processes. Therefore, an immediate force feedback is crucial to correlate the applied forces and the triggered intracellular effects, such as the adaption of growth, changes in material properties, or intracellular signaling cascades that are set in motion. Until now, direct measurements of forces exerted by tip-growing cells have only been performed on fungal hyphae, where the measurement was conducted with a silicon bridge strain gauge (Johns *et al*., 1999; Money, 2007). However, the low-resolution force of 1 µN, the manual calibration, and the inadaptability of the orientation of the sensor with respect to the hyphae, have hindered the broad use of this technique.

Merging the flexibility of microfluidic design with the direct and immediate force feedback of a MEMS-based capacitive force sensor (Enikov and Nelson, 2000; Sun and Nelson, 2007) opens up new ways to investigate the effect of forces and stress on the growth and physiology of tip-growing cells. In this study, an *in vitro* environment was designed by combining microfluidics and MEMS force-sensing to mimic the PT’s journey from stigma to ovary while simultaneously assessing the arising forces and observe the changes in the PT’s growth behavior. The perceptive force, defined as the force at which the PTs react to a mechanical barrier by changing their growth direction, has been determined in two different species, *Lilium longiflorum* (lily) and *Arabidopsis thaliana*. Furthermore, the penetrative force of the PTs, when they squeeze through a microgap, was quantified and correlated to geometrical factors and turgor pressure.

## Methods

### Control and Data Acquisition

The lateral force sensor (FT-S1000-LAT, FemtoTools AG) is able to measure a force range of ±1000 µN with a resolution of 0.05 µN at 10 Hz. The force sensor was mounted on an aluminum arm on a piezoelectric xyz positioner for precise threedimensional control. The experiments were recorded with a 40x objective lens (LUCPlanFL N 40x, Olympus) mounted on an inverted microscope (IX71, Olympus) using a CCD camera (Orca-D2, Hamamatsu). An xy stage (M-687.UO, Physik Instrumente (PI) GmbH & Co.) integrated onto the microscope was actuated by a piezomotor controller (C-867.262, PI GmbH & Co.) and used to position the sample in the field of view. Data acquisition and the control of the xy stage and the xyz positioner were implemented in LabVIEW™ (National Instruments (NI)). Data analysis was performed in MATLAB (MathWorks).

### Pollen Tube Preparation

Anthers of *Lilium longiflorum* were placed in Eppendorf tubes, and stored at −80 °C. To germinate the PTs, pollen grains were rehydrated in humid chambers for 30 min at room temperature, after which they were mixed with PT growth medium (PTGM) and filled into the LOC device. PTs were grown in solidified growth medium. Lily PTGM is: 160 µM H_3_BO_3_, 130 µM Ca(NO_3_)_2_, 1 mM KNO_3_, 5 mM MES and 10 % sucrose at a pH of 5.5. To solidify the medium, we added 1 % ultra-low-gelling agarose (A2576, Sigma-Aldrich). Solid PTGM was stored at 4 °C, re-melted at 63 °C, and equilibrated at 23 °C before mixing with rehydrated pollen grains and loading into the LOC device. To solidify the agarose within the microchannels, the LOC was cooled at 4 °C for 4 min. Excess agarose in front of the channels was removed with a wooden toothpick. This is important to avoid interference of agarose with the force sensor. The reason for using solidified growth medium was to simulate the mechanical resistance of the transmitting tissue in plants. As a positive side effect, the pollen in the agarose solution showed a more stable growth pattern over a longer time periods compared to those growing in liquid PTGM. The glass slides in front of the LOC where the PTs emerge were treated with poly-*L*-lysine (P4832-50ML, Sigma-Aldrich), forming a monolayer to improve the adhesion of the PTs to the glass slide.

Flowers of *Arabidopsis thaliana*, accession Col-0, were collected and incubated for 30 min at 22 °C in a moisture chamber. PTs were grown in solidified PTGM (1.6 mM H_3_BO_3_, 5 mM CaCl_2_, 5 mM KCl, 1 mM MgSO_4_, and 10 % sucrose at a pH of 7.5). The solidification of agarose and LOC loading followed the same procedure as described for lily PTs.

## Results

### Measuring the forces that are involved when pollen tubes are confronted with obstacles

To simulate the natural environment of the pollen tube *in vitro*, we designed a system that forces the PTs to grow against a mechanical barrier or to squeeze through narrow gaps, while we simultaneously recorded the generated forces in real-time. The system combines a cellular force microscope (CFM) (Vogler *et al*., 2013; Felekis *et al*., 2011; Felekis *et al*., 2014; Felekis *et al*., 2015), equipped with a commercially available lateral force sensor, with a LOC for high-throughput pollen germination and guided parallelized PT growth, adapted from previous work by Shamsudhin and colleagues (2016) (Fig. 1A). The LOC provides a reservoir in which the pollen grains germinate. After germination, the PTs grow into parallel microchannels with square cross-sectional areas of 25×25 µm (Fig. 1B). The capacitive MEMS sensor offers a high-resolution force-sensing plate of 50×50 µm (Fig. 1C). The force plate is placed at the end of the microchannels and acts as an obstacle with integrated force feedback (Fig. 1D). The distance between the force plate and the microchannel can be adjusted to form a narrow gap, mimicking the small cavities in the apoplast between cells along the PT’s path to the ovule.

**Fig 1.**
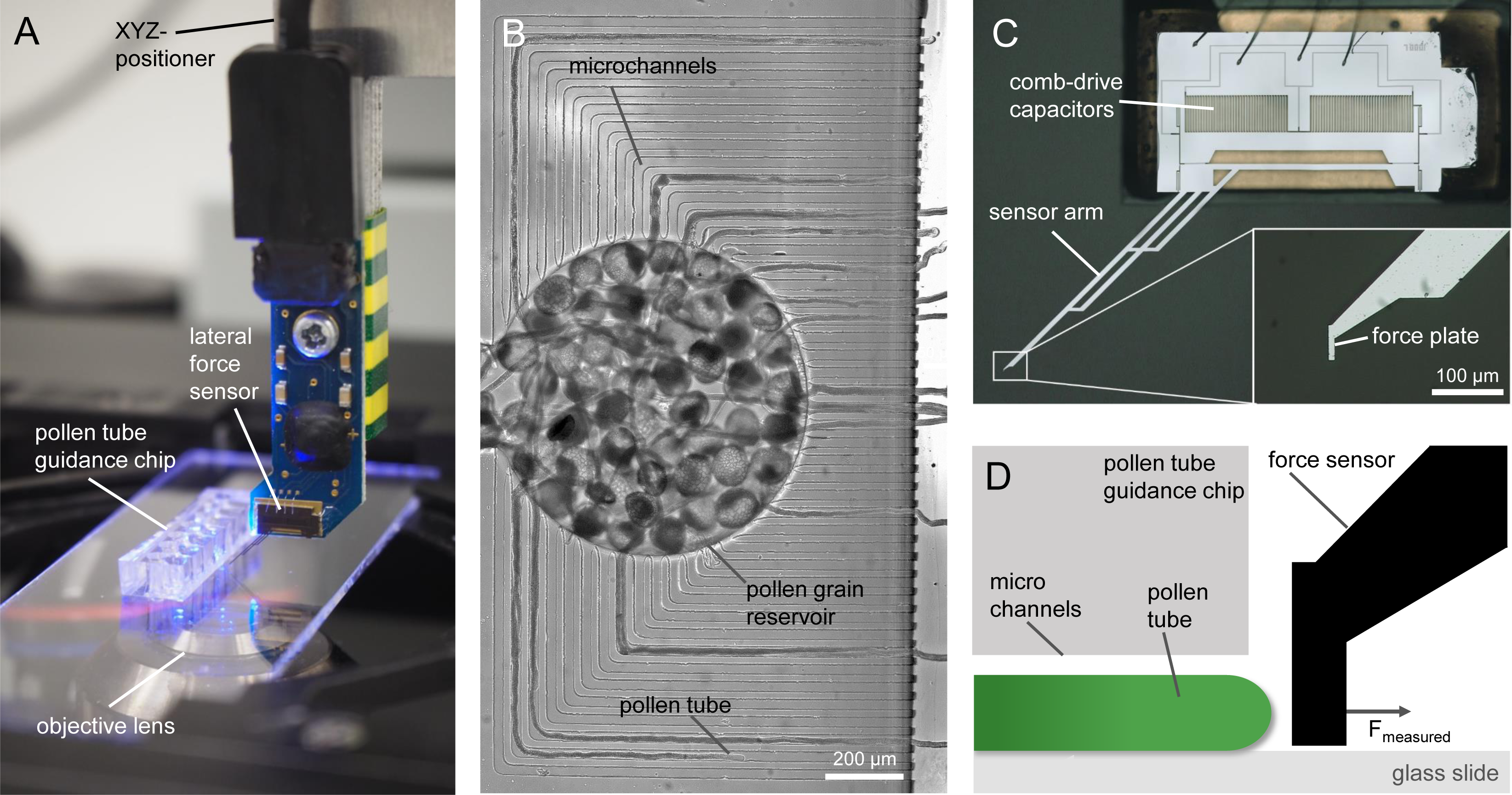
System configuration to simulate the natural environment of the pollen tube in vitro with integrated force feedback. (A) The experimental system combines a LOC device with a MEMS-based force sensor. (B) An optical image of a LOC loaded with pollen grains in the inner reservoir; the PTs grow in parallel in the same focal plane through the microchannels. (C) Capacitive force sensor measuring lateral forces at the force-sensing plate. (D) Schematic side-view of the experimental setup where the force-sensing plate is placed in front of the PTs emerging from the microchannels.

Mounted on an inverted microscope, this design permits direct force measurements and high-resolution imaging to capture the real-time response of PTs when interacting with mechanical obstacles.

### Pollen tubes perceive the force exerted by obstacles and react with a change in growth behavior

The ability to sense mechanical stress or forces is crucial for cells to adapt their growth and shape according to changing environmental conditions. PTs with their fast growth rates and their invasive behavior must rapidly adapt their growth according to external mechanical stimuli. The interaction of the growing PTs with physical barriers and the forces generated reveal a tactile perception in lily PTs.

In our experiments, the PTs were guided along parallel microchannels to achieve unidirectional growth and to ensure a perpendicular contact with the force-sensing plate placed at a distance larger than the PT diameter from the channel exit. This gap allowed the PT to change growth direction without restrictions from the LOC channels. Using the force-sensing plate as a mechanical obstacle offered real-time force feedback, which allowed us to measure the force parallel to the growth direction of the PTs. After the initial contact of the PTs with the sensor plate, a sharp force increase was observed (Fig. 2B i) while the apical region was symmetrically compressed (Fig. 2A i-ii). This passive deformation was observed up to a force with a mean value of 9.6 µN and a standard deviation of 1.6 µN in lily (Fig. 2B ii and 2C). After that point, the force remained constant, indicating that the PT stopped growing (Fig. 2B ii-iii). After a short lag phase (typically a few seconds), the PT resumed growth, which was accompanied by a change in growth direction, visible as a small bulb perpendicular to the original growth axis (Fig. 2A iii). At the last stage, the force remained at a constant level (Fig. 2B iv) while the PT tip grew along the force plate (Fig. 2A iv).

**Fig 2.**
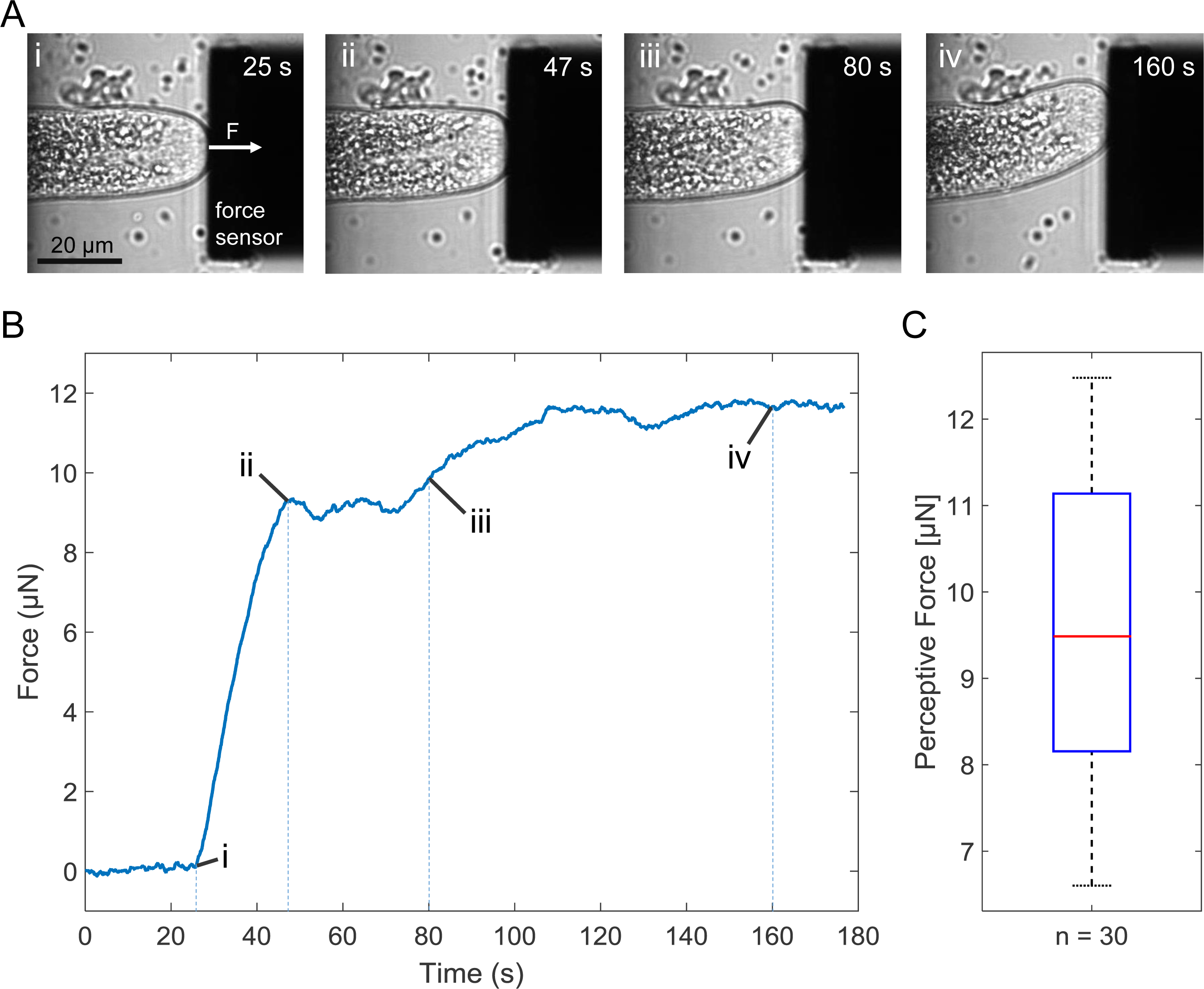
Force perception of lily pollen tubes when confronted with an obstacle measured by a MEMS force sensor. (A) The PTs adjust their apical growth direction in response to external mechanical cues: (i) initial contact, (ii) passive deformation, (iii) adjustment of apical growth direction, and (iv) bending of the shank. (B) Forces generated between PT tip and force sensing plate during the interaction: (i) initial contact, (ii) perceptive force, (iii-iv) stable force. (C) Descriptive statistics of the perceptive force in lily PTs. Red line indicates the median, blue box indicates the 25^*th*^ and 75^*th*^ percentile and the whiskers extend to the most extreme data points not considered outliers.

The observed growth arrest indicates that lily PTs can sense physical obstacles and react to this external mechanical stimulus by adjusting their growth direction in order to avoid obstacles. Since this critical force threshold marks the moment that the PT perceives and responses to the force, we refer to it as the perceptive force (Fig. 2C).

### Force perception is not species-specific

Lily PTs have a comparably large diameter and are easy to manipulate. Their disadvantage is that genetic tools, such as mutant or transgenic lines, do not exist, making it difficult to examine the physiological and molecular basis of force perception. However, all these tools are available in Arabidopsis, another model species in plant biology. Therefore, we repeated the experiments using Arabidopsis PTs to test whether the observed force sensing behavior was general or species-specific. The observed change in growth (Fig. 3A) together with the force pattern (Fig. 3B) revealed that Arabidopsis PTs behave analogously to lily PTs when confronted with a physical barrier. In the case of Arabidopsis, however, a perceptive force of 3.0 µN with a standard deviation of 0.6 µN was measured (Fig. 3C), which is considerably lower than the one measured for lily PTs. The subsequent change in growth direction was more prominent in Arabidopsis, leading to a sharp, almost perpendicular bending after interacting with the force-sensing plate (Fig. 3A iv). The results of the comparison between lily and Arabidopsis suggest that force sensing in PTs is a conserved mechanism, which is common to mono- and dicotyledonous plant species.

**Fig 3.**
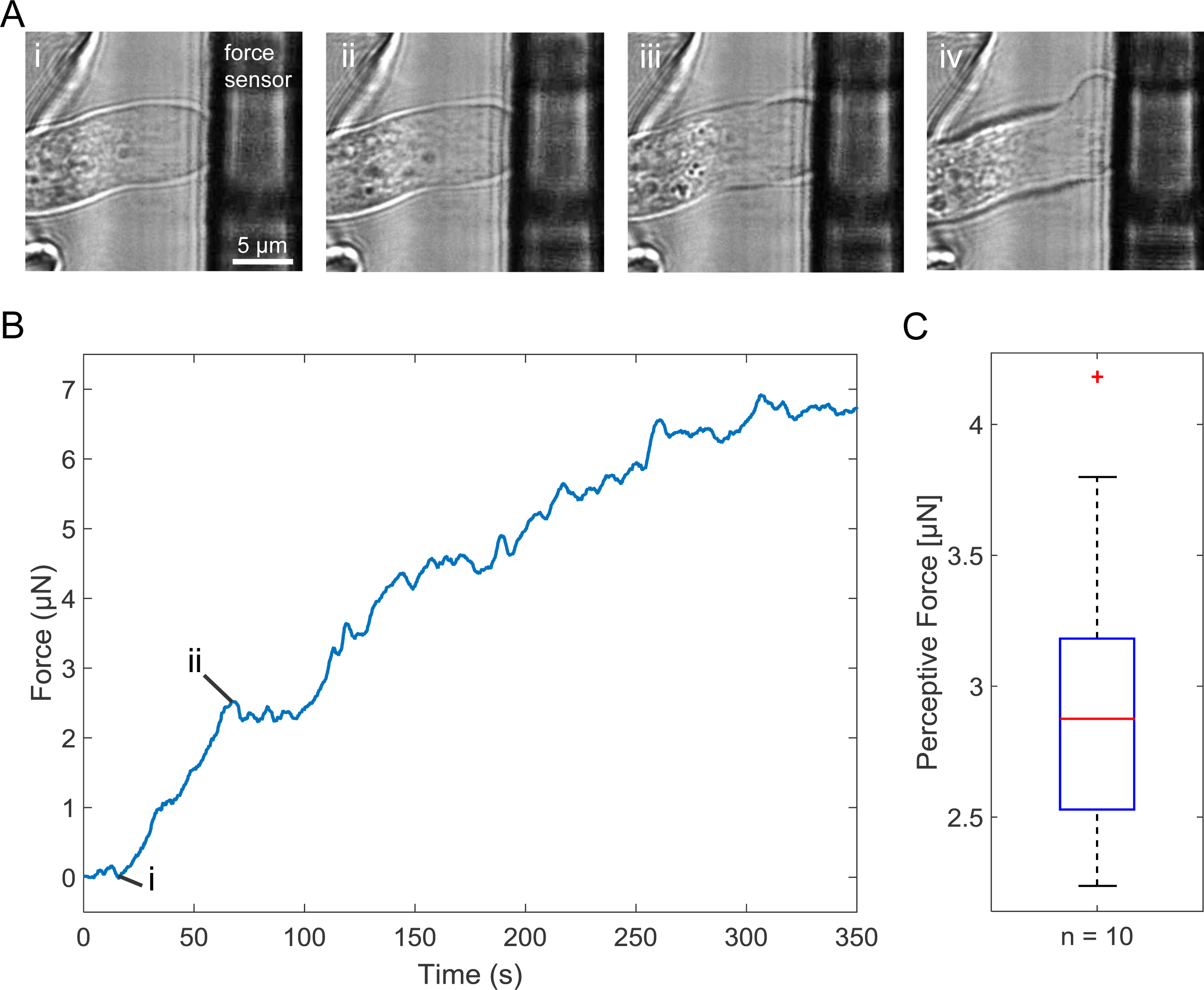
Force perception in Arabidopsis pollen tubes. (A) Similar to the lily PTs, Arabidopsis PTs adjust their apical growth direction in response to external mechanical cues: (i) initial contact, (ii) passive deformation, (iii-iv) adjustment of apical growth direction. (B) Forces generated between the PT tip and the force plate during the interaction. (i) initial contact and (ii) perceptive force. (C) Descriptive statistics of the perceptive force in Arabidopsis PTs. Red line indicates the median, blue box indicates the 25^*th*^ and 75^*th*^ percentile, red cross indicates outliers and the whiskers extend to the most extreme data points not considered outliers.

### Invasive Growth: Penetration follows Perception

Apart from sensing and reacting to external mechanical obstacles, PTs can exert forces on their environment. This is crucial for invading cells in surrounding tissue (i.e. stigmatic papillar cells) or squeezing between transmitting tract cells, and eventually delivering the sperm cells to the ovules. In our experiment, the force sensing plate was placed close to the exit of the microchannels forming a narrow gap (smaller than the PT diameter), through which the PT could penetrate (Fig. 4A). The gap simulated the path of the PT through the apoplast growing within or between cells, and the penetration of the ovule through the micropyle and the filiform apparatus.

**Fig 4.**
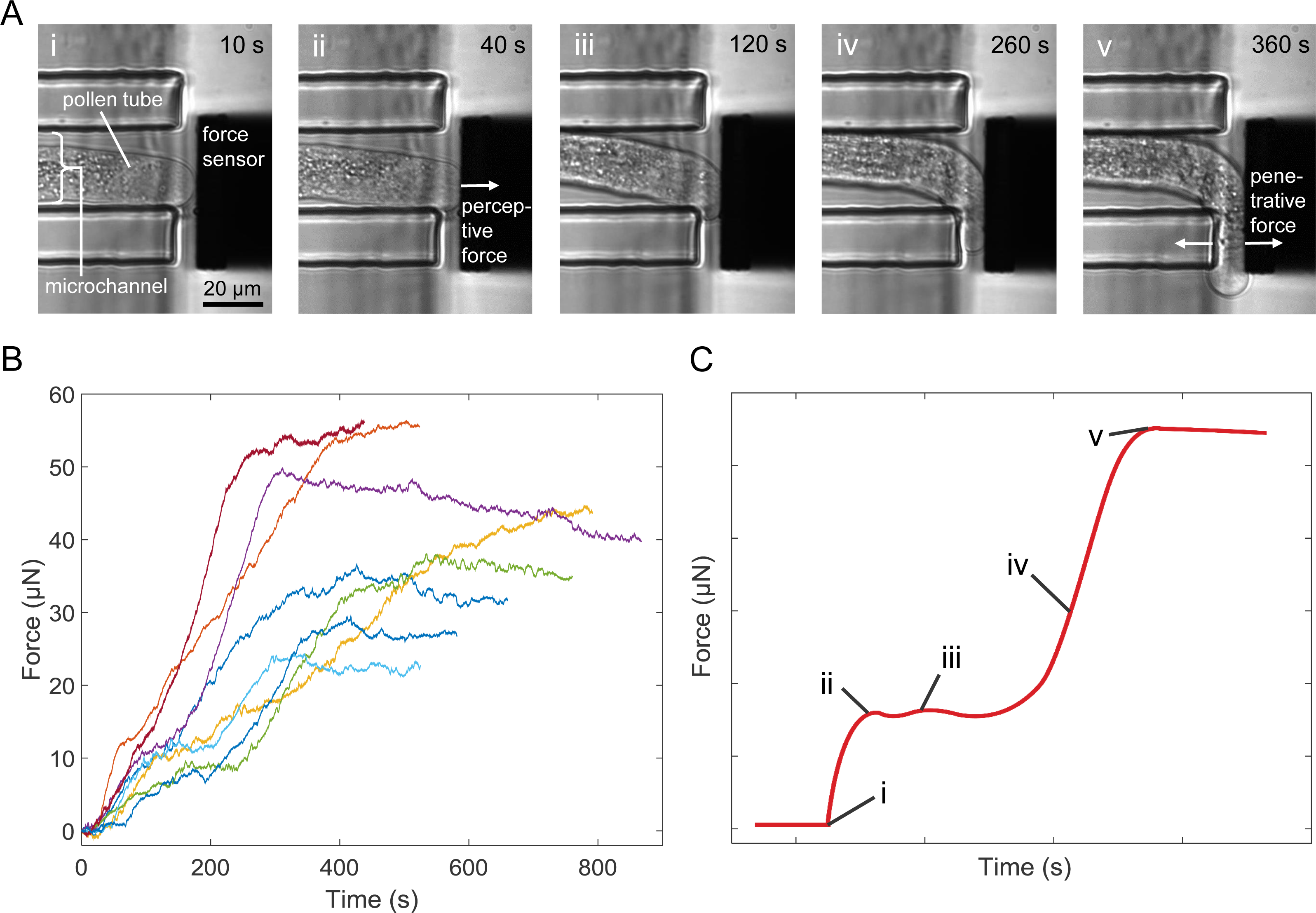
Forces exerted by lily pollen tubes growing through a small gap (A) Five different stages of the PT’s interaction to sense and overcome a barrier by penetrating through a narrow gap. (B) Force curves of individual PTs when confronted with a mechanical barrier (force sensor). All the tested PTs show a similar response when interacting with a mechanical obstacle. (C) A schematic force curve representing the akin force curves and highlighting the different stages.

Lily PTs showed a clear sequence of behaviors when squeezing through the artificially created barrier. This sequence consisted of five stages corresponding to initial contact, perception, adaptation, penetration, and emergence. The first two stages (Fig. 4A i-ii) were equivalent to the interactions discussed in the previous section. After recognizing the mechanical barrier, the PTs adjusted their growth direction and, when confronted with a narrow opening, anchored themselves with a small protrusion within the gap (Fig. 4A iii). If the apex was already close to the small opening this phase was very short, but occasionally the PT first bent into several directions until it located a suitable interstice. Once detected, the PT squeezed into the slim lacuna and slightly widened it by the forces it generated (Fig. 4A iv). At last, the PT emerged from the channel and continued to grow while returning to its original shape (Fig. 4A v).

The measured force was correlated with the individual phases during the interaction. In the experiment, we observed that different PTs showed similar force patterns (Fig. 4B) and a representative schematic force curve is illustrated in Fig. 4C, where the five penetration stages are highlighted. The force sharply increased after the PT contacted the force plate (Fig. 4C i). After perceiving the mechanical obstacle, the force stopped increasing and formed a small plateau in the force curve (Fig. 4C ii). When the PT adapted its growth direction and sought a way to overcome the barrier, the forces varied around a constant value (Fig. 4C iii). With the chasm found, the slope of the force curve increased while the apex entered the small gap. Once the whole apex had completely entered the constriction, an almost linear increase of the force with time was observed (Fig. 4C iv). When the PT emerged from the gap, the force remained constant (Fig. 4C v).

The penetrative force, generated at the flank of the PT when squeezing through the gap, was obtained by subtracting the force values prior to the penetration (i.e. the perceptive force) from the force measured after emerging, and was in the range of several tens of µN up to 84.91 µN, with a mean value of 29 µN and a standard deviation of 18 µN (Fig. 5A). The large variation of these values arose due to dependencies of the measured maximal forces on geometrical factors of the gap and the PT itself.

**Fig 5.**
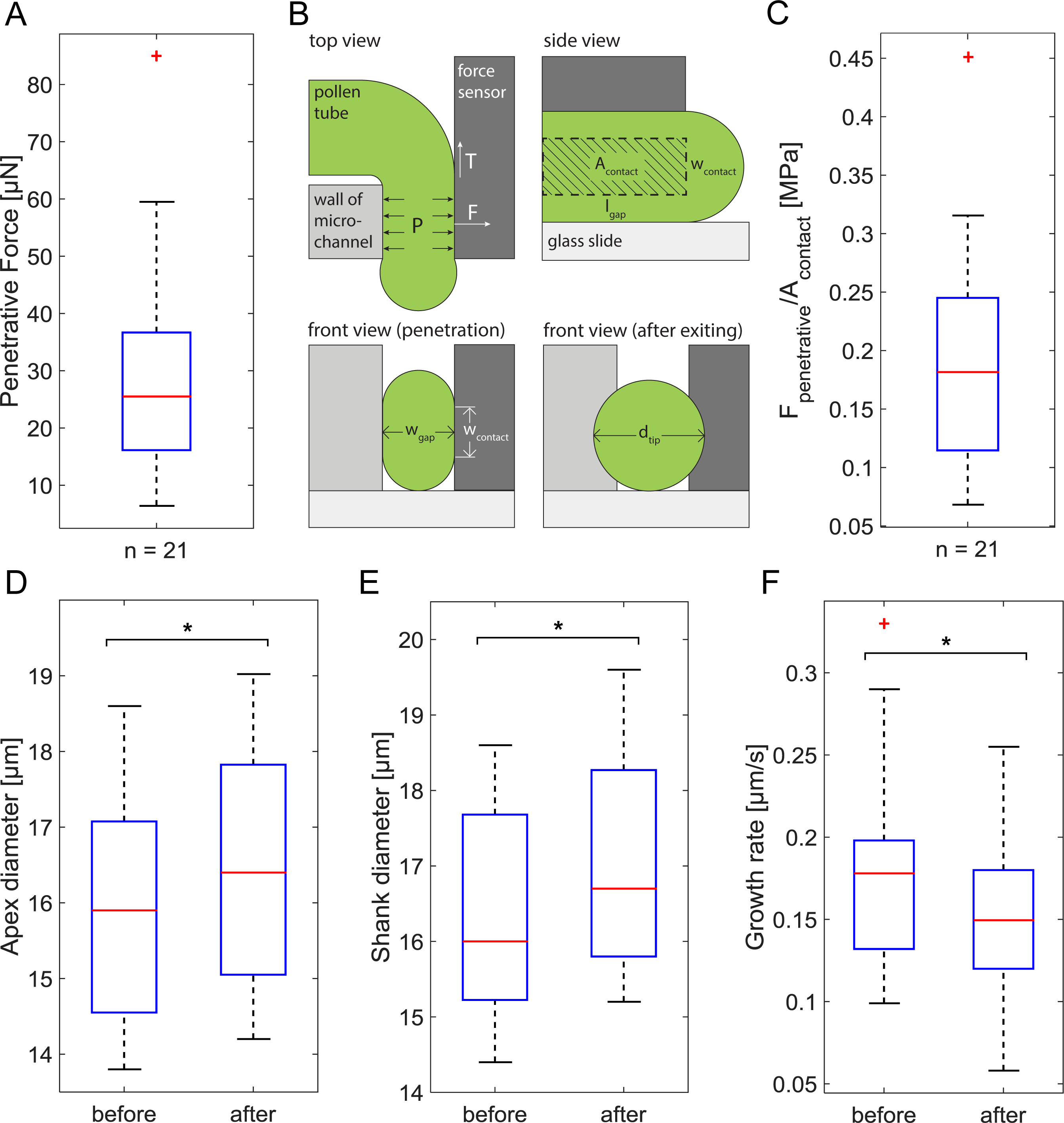
Descriptive statistics of generated forces, turgor pressure and change in pollen tube growth rate and diameter before and after penetration. (A) Penetrative force generated by lily pollen tubes. (B) Geometrical feature of the gap and the PT allowing an approximation of the contact area. (C) Ratio between the penetrative force and the contact area yields the turgor pressure of the PTs after they emerged. (D) Apex diameter, (E) shank diameter, and (F) growth rate of the PTs before and after penetration through the gap. (*) indicates a significant difference verified by a two-tailed, paired t-test and *p* < 0.01. Boxplots: Red line indicates the median, blue box indicates the 25^*th*^ and 75^*th*^ percentile, red cross indicates outliers, and the whiskers extend to the most extreme data points not considered outliers.

The penetrative force (*F_p_*) is given by the contact area *A* multiplied with the turgor pressure *P* at the time when the PTs emerge from the gap. Since the cell wall that is in contact with the force plate runs parallel to it, its tension *T* does not contribute to the measured forces (Fig. 5B). The width and length of the gap, as well as the diameter of the PT after emerging can be extracted from the optical recordings. The contact area of the PT with the sensor can be estimated using these values. Assuming that the cell wall does not expand, we can equate the perimeter U of the apex after emerging with the one during the penetration,

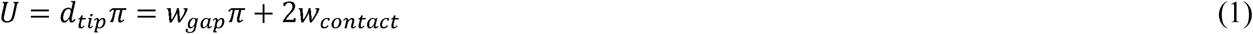

The contact area A is then determined by

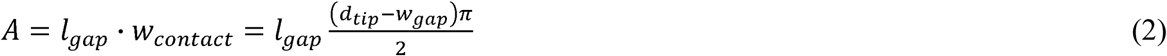

where d_tip_ is the PT tip diameter, w_gap_ the gap width, l_gap_ the gap length, and wcontact the contact width between the PT and sensor plate (Fig. 5B).

The ratio between penetrative force *F_p_* and contact area *A* yields the turgor pressure *P* (Fig. 5C) of the PT when it grows out from the channel according to

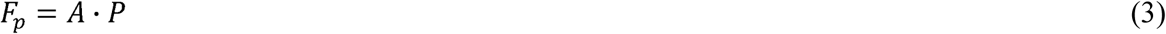

A turgor pressure of (0.19 ± 0.09) MPa was derived from our measurements, which is consistent with the reported values of 0.1 MPa to 0.4 MPa for lily (Benkert *et al*., 1997).

PT dimensions changed after emerging from the narrow openings. The apex expanded back to its original round shape after emergence, but with a significantly increased diameter compared to its initial state (Fig. 5D). The diameter of the shank, measured at a fixed position in the microchannel, increased as well. Since cell wall composition in the shank is not modified, this implies an increase in turgor pressure during this physical interaction (Fig. 5E). Interestingly, the growth rate of the PTs was smaller after emerging from the gap than before entering it (Fig. 5F). We speculate that the cytoskeletal organization could be disturbed through the geometrical deformation, leading to problems with the transport of new cell wall material to the PT tip. Alternatively, the demand for cell wall material could be increased due to the increased diameter of the apical region.

The penetrative force depends mainly on the geometry of the gap. An upper bound of these forces is only given when the gap is too small for the PT to invade or if the gap is too long for the PT to penetrate. The force is created by the deformation of the PT and the turgor pressure. Additionally, the PT is able to increase the turgor pressure in order to generate a larger force.

## Discussion

The experimental system combines a LOC device and a MEMS force sensor to create physical obstacles with an integrated force feedback. The growth of tip growing single cells, e.g. PTs, is directionally guided onto a defined obstacle with adjustable gap sizes. This allows the study of interactions between tip growing cells and mechanical barriers, thereby assessing the influence of external mechanical stimuli onto their perceptive behavior and response. The capability to directly measure the forces arising between the cell and the sensor plate can immediately correlate force changes with the corresponding growth behavior and adaption.

We focused our experiments on PTs, but the configuration can be adapted to characterize other tip growing cells, such as root hairs, moss protonemata, or fungal hyphae. PTs are of particular interest because of their remarkable capability to precisely navigate to the ovule by covering tremendous distances (e.g. 300 mm in maize), while invading and penetrating tissues and cells. Their growth is guided by biochemical, electrical, and mechanical cues, but it is largely unknown how these cues are internally processed.

Our system is designed to study how PTs perceive mechanical cues and adapt their growth behavior to navigate around obstacles and squeeze through narrow openings. Their reaction to external mechanical stress as well as the forces they exert on their environment were monitored. A perceptive force of 9.6 µN in lily and 3.0 µN in Arabidopsis PTs was observed. After the perceptive force was reached, the PTs sensed the obstacle and adjusted their apical growth direction in order to avoid it. This suggests the presence of mechanosensitive receptors in the apical region.

PTs were able to penetrate through tiny gaps by first anchoring themselves with a small protrusion. The smaller and longer the gap the PT had to penetrate through, the larger the observed force. Penetrative forces of up to 84.9 µN were measured. An estimation of the contact area allowed us to calculate the pressure the PT exerted on the force plate, which corresponded to the turgor pressure of the PT at the moment when it emerged from the gap. Hence, the force solely depended on the turgor pressure and the PT’s ability to modify it. The increase in turgor pressure after penetration, observed by an increase in the diameter of the shank, facilitates the invasion of tissues and cells since larger forces can be generated.

The design of the LOC facilitates microscopy by confining the PTs to a single focal plane. Hence, intracellular activities can be investigated during the PT–force sensor interactions, for example by observing the effects of mechanical stress on physiological parameters using fluorescent biosensor molecules. Therefore, our system will be instrumental in identifying new molecules and pathways that are involved in mechanotransduction.

## Supplementary Data

Video S1. Force perception of a lily pollen tube when confronted with an obstacle measured by a MEMS force sensor.

Video S2. Forces exerted by a lily pollen tube growing through a small gap.

## Acknowledgements

This work was financed through ETH Zürich and the University of Zürich, and a Research and Technology Development (RTD) project (MecanX - Understanding Physics of Plant Growth) of SystemsX.ch, the Swiss Initiative in Systems Biology (to UG and BJN).

## Author contributions

UG and BJN initiated and supervised the project, and raised funding. JTB and HV conceived, conducted, and analyzed the experiments, NFL designed and fabricated the LOC device. JTB and HV wrote the manuscript. All authors reviewed and commented on the manuscript.

## References

Agudelo CG, Sanati Nezhad A, Ghanbari M, Naghavi M, Packirisamy M, Geitmann A. 2013. TipChip: a modular, MEMS-based platform for experimentation and phenotyping of tip-growing cells. The Plant Journal 73, 1057–1068.

Benkert R, Obermeyer G, Bentrup F-W. 1997. The turgor pressure of growing lily pollen tubes. Protoplasma 198, 1–8.

Borg M, Brownfield L, Twell D. 2009. Male gametophyte development: a molecular perspective. Journal of Experimental Botany 60, 1465–1478.

Braam J. 2004. In touch: plant responses to mechanical stimuli. New Phytologist 165, 373-389.

Chebli Y, Geitmann A. 2007. Mechanical principles governing pollen tube growth. Functional Plant Science and Biotechnology 1, 232–245.

Chehab EW, Eich E, Braam J. 2009. Thigmomorphogenesis: a complex plant response to mechano-stimulation. Journal of Experimental Botany 60, 43–56.

Darwin C, Darwin F. 1880. The power of movement in plants. Appleton.

Ellinger D, Voigt C. 2014. Callose biosynthesis in Arabidopsis with a focus on pathogen response: what we have learned within the last decade. Annals of Botany 114, 1349–1358.

Enikov ET, Nelson BJ. 2000. Three-dimensional microfabrication for a multi-degree-of-freedom capacitive force sensor using fibre-chip coupling. Journal of Micromechanics and Microengineering 10, 492–497.

Felekis D, Muntwyler S, Vogler H, Beyeler F, Grossniklaus U, Nelson BJ. (2011). Quantifying growth mechanics of living, growing plant cells in situ using microrobotics. Micro & Nano Letters 6, 311-316.

Felekis D, Vogler H, Mecja G, Muntwyler S, Nestorova A, Huang T, Sakar MS, Grossniklaus U, Nelson BJ. 2015. Real-time automated characterization of 3D morphology and mechanics of developing plant cells. The International Journal of Robotics Research 34, 1136–1146.

Felekis D, Vogler H, Mecja G, Muntwyler S, Sakar MS, Grossniklaus U, Nelson BJ. 2014. High-throughput analysis of the morphology and mechanics of tip growing cells using a microrobotic platform. IEEE International Conference on Intelligent Robots and Systems, 3955–3960.

Haley A, Russell AJ, Wood N, Allan AC, Knight M, Campbell AK, Trewavas AJ. 1995. Effects of mechanical signaling on plant cell cytosolic calcium. Proceedings of the National Academy of Sciences of the United States of America 92, 4124–4128.

Hamant O, Heisler M, Jönsson H, Krupinski P, Uyttewaal M, Bokov P, Corson F, Sahlin P, Boudaoud A, Meyerowitz E, Couder Y, Traas J. 2008. Developmental patterning by mechanical signals in Arabidopsis. Science 322, 1650–1655.

Haswell ES, Peyronnet R, Barbier-Brygoo H, Meyerowitz EM, Frachisse J-M. 2008. Two MscS homologs provide mechanosensitive channel activities in the Arabidopsis root. Current Biology 18, 730–734.

Heisler M, Hamant O, Krupinski P, Uyttewaal M, Ohno C, Jönsson H, Traas J, Meyerowitz E. 2010. Alignment between PIN1 polarity and microtubule orientation in the shoot apical meristem reveals a tight coupling between morphogenesis and auxin transport. PLoS Biology 8, e1000516.

Higashiyama T, Kuroiwa H, Kuroiwa T. 2003. Pollen-tube guidance: beacons from the female gametophyte. Current Opinion in Plant Biology 6, 36–41.

Higashiyama T, Yang W-C. 2017. Gametophytic pollen tube guidance: attractant peptides, gametic controls, and receptors. Plant Physiology 173, 112–121.

Horade M, Kanaoka MM, Kuzuya M, Higashiyama T, Kaji N. 2013. A microfluidic device for quantitative analysis of chemoattraction in plants. RSC Advances 3, 22301-22307.

Hu C, Munglani G, Vogler H, Fabrice TN, Shamsudhin N, Wittel FK, Ringli C, Grossniklaus U, Herrmann HJ, Nelson BJ. 2017a. Characterization of size-dependent mechanical properties of tip-growing cells using a lab-on-chip device. Lab on a Chip 17, 82–90.

Hu C, Vogler H, Aellen M, Shamsudhin N, Jang B, Burri JT, Läubli N, Grossniklaus U, Pané S, Nelson BJ. 2017b. High precision, localized proton gradients and fluxes generated by a microelectrode device induce differential growth behaviors of pollen tubes. Lab on a Chip 17, 671–680.

Hülskamp M, Schneitz K, Pruitt R. 1995. Genetic evidence for a long-range activity that directs pollen tube guidance in Arabidopsis. The Plant Cell 7, 57–64.

Jaffe MJ. 1973. Thigmomorphogenesis: the response of plant growth and development to mechanical stimulation. Planta 114, 143–157.

Jaffe MJ, Forbes S. 1993. Thigmomorphogenesis: the effect of mechanical perturbation on plants. Plant Growth Regulation 12, 313–324.

John S, Davis CM, Money NP. 1999. Pulses in turgor pressure and water potential: resolving the mechanics of hyphal growth. Microbiological Research 154, 225–231.

Kesten C, Menna A, Sànchez-Rodrìguez C. 2017. Regulation of cellulose synthesis in response to stress. Current Opinion in Plant Biology 40, 106–113.

Legué V, Blancaflor E, Wymer C, Perbal G, Fantin D, Gilroy S. 1997. Cytoplasmic free Ca^2+^ in Arabidopsis roots changes in response to touch but not gravity. Plant Physiology 114, 789–800.

Lintilhac PM, Vesecky TB. 1981. Mechanical stress and cell wall orientation in plants. II. The application of controlled directional stress to growing plants; with a discussion on the nature of the wound reaction. American Journal of Botany 68, 1222–1230.

Lynch TM, Lintilhac PM. 1997. Mechanical signals in plant development: a new method for single cell studies. Developmental Biology 181, 246–256.

Mollet JC, Park SY, Nothnagel EA, Lord, EM. 2000. A lily stylar pectin is necessary for pollen tube adhesion to an in vitro stylar matrix. The Plant Cell 12, 1737–1749.

Money NP. 2007. Biomechanics of invasive hyphal growth. In: Howard RJ, Gow NAR., eds. Biology of the fungal cell. Heidelberg: Springer, 237–249.

Monshausen GB, Haswell ES. 2013. A force of nature: molecular mechanisms of mechanoperception in plants. Journal of Experimental Botany 64, 4663–4680.

Prado AM, Porterfield DM, Feijó, JA. 2004. Nitric oxide is involved in growth regulation and re-orientation of pollen tubes. Development 131, 2707–2714.

Qu LJ, Li L, Lan Z, Dresselhaus T. 2015. Peptide signaling during the pollen tube journey and double fertilization. Journal of Experimental Botany 66, 5139–5150.

Ray SM, Park SS, Ray A. 1997. Pollen tube guidance by the female gametophyte. Development 124, 2489–2498.

Sanati Nezhad A, Naghavi M, Packirisamy M, Bhat R, Geitmann A. 2013. Quantification of cellular penetrative forces using lab-on-a-chip technology and finite element modeling. Proceedings of the National Academy of Sciences 110, 8093-8098.

Sanati Nezhad A, Packirisamy M, Geitmann A. 2014. Dynamic, high precision targeting of growth modulating agents is able to trigger pollen tube growth reorientation. The Plant Journal 80, 185–195.

Shamsudhin N, Laeubli N, Atakan HB, Vogler H, Hu C, Haeberle W, Sebastian A, Grossniklaus U, Nelson BJ. 2016. Massively parallelized pollen tube guidance and mechanical measurements on a lab-on-a-chip platform. PLoS ONE 11, e0168138.

Sun Y, Nelson BJ. 2007. MEMS capacitive force sensors for cellular and flight biomechanics. Biomedical Materials 2, 16-22.

Vogler H, Draeger C, Weber A, Felekis D, Eichenberger C, Routier-Kierzkowska A-L, Boisson-Dernier A, Ringli C, Nelson BJ, Smith RS, Grossniklaus U. 2013. The pollen tube: a soft shell with a hard core. The Plant Journal 73, 617–627.

Wolters-Arts M, Lush WM, Mariani C. 1998. Lipids are required for directional pollen-tube growth. Nature 392, 818–821.

Wymer CL, Wymer SA, Cosgrove DJ, Cyr RJ. 1996. Plant cell growth responds to external forces and the response requires lntact microtubules. Plant Physiology 110, 425–430.

Yahraus T, Chandra S, Legendre L, Low PS. 1995. Evidence for a mechanically induced oxidative burst. Plant Physiology 109, 1259–1266.

Yanagisawa N, Sugimoto N, Arata H, Higashiyama T. 2017. Capability of tip-growing plant cells to penetrate into extremely narrow gaps. Scientific Reports 7, 1403.

